# Direct Observation of Heterogeneous Formation of Amyloid Spherulites in Real-time by Super-resolution Microscopy

**DOI:** 10.1101/2021.08.20.457097

**Authors:** Min Zhang, Henrik D. Pinholt, Xin Zhou, Søren S.-R. Bohr, Luca Banetta, Alessio Zaccone, Vito Foderà, Nikos S. Hatzakis

## Abstract

The misfolding of proteins and their aggregation in the form of fibrils or amyloid-like spherulites are involved in a spectrum of pathological abnormalities. Our current understanding of protein amyloid aggregation mechanisms has primarily relied on the use of spectrometric methods to determine the average growth rates and diffraction limited microscopes with low temporal resolution to observe the large-scale morphologies of intermediates. We developed a REal-time kinetics via binding and Photobleaching LOcalisation Microscopy (**REPLOM**) super-resolution method to directly observe and quantify the existence and abundance of diverse aggregate morphologies below the diffraction limit and extract their heterogeneous growth kinetics. Our results revealed that even the growth of a microscopically identical aggregates, e.g. amyloid spherulites, may follow distinct pathways. Specifically, spherulites do not exclusively grow isotropically but, surprisingly, may also grow anisotropically, following similar pathways as reported for minerals and polymers. Combining our technique with machine learning approaches, we associated growth rates to specific morphological transitions and provided energy barriers and the energy landscape at the level of single aggregate morphology. Our unifying framework for the detection and analysis of spherulite growth can be extended to other self-assembled systems characterized by a high degree of heterogeneity, disentangling the broad spectrum of diverse morphologies at the single-molecule level.

## Introduction

Protein misfolding and aggregation in the form of fibrils or spherulites are a hallmark of a number of devastating conditions, such as Alzheimer’s and Parkinson’s disease (Chiti & Dobson, 2006; Exley et al., 2010; House, Jones, & Exley, 2011). Indeed, elongated protein aggregates, known as amyloid-like fibrils, are a characteristic of these diseases and, in the last two decades, deciphering the key steps of their formation has been the main focus of the amyloid research community (Nielsen et al., 2001; Pinotsi, Buell, Dobson, Schierle, & Kaminski, 2013; Zimmermann et al., 2021). However, a “one-size-fits-all” approach to describing amyloid-forming systems will not be successful. Other amyloid-like species (named *superstructures*) may occur that exhibit significantly different β-sheet packing compared to fibrils (Vito Foderà et al., 2014; Vetri & Foderà, 2015). These include amyloid-like spherical aggregates, or spherulites, that range from a few micrometers to several millimeters in diameter and can form both *in vivo* and *in vitro* (Exley et al., 2010; Mark R. H. Krebs et al., 2004; Vetri & Foderà, 2015). These aggregates are characterized by a fascinating core-shell morphology and seem to be the result of a general self-assembly process that is common to metal alloys (Lu, Goh, Li, & Ng, 1999), minerals (Heaney & Davis, 1995), and polymers (Hosier, Bassett, & Vaughan, 2000; Kajioka, Hikosaka, Taguchi, & Toda, 2008). Protein amyloid spherulites are hallmarks of disease states. Specifically, deposition in brain tissues of amyloid spherulites consisting of Aβ peptide has been found in connection with the onset and progression of Alzheimer disease (Exley et al., 2010; House et al., 2011). In addition, they may also present opportunities to develop advanced materials for drug delivery (Jiang et al.). While we have a solid understanding of the fibrillar growth kinetics (Garcia, Cohen, Dobson, & Knowles, 2014), the mechanisms of the formation and growth of spherulites is still out of reach (Ban et al., 2006; Domike & Donald, 2007; M. R. H. Krebs, Bromley, Rogers, & Donald, 2005).

Current models of the mechanism of protein spherulite formation primarily rely on spectrometric evidence for their average growth rates (Domike & Donald, 2009; M. R. H. Krebs et al., 2005), low temporal resolution recordings of growth intermediates (Ban et al., 2006; Yagi, Ban, Morigaki, Naiki, & Goto, 2007) and observations of the final structures via microscopy techniques (Mark R. H. Krebs et al., 2004; Toprakcioglu, Challa, Xu, & Knowles, 2019). The evidence has resulted in a hypothesis that spherulites are core-shell structures in which fibril-like filaments isotropically and radially grow around a dense core (Mark R. H. Krebs et al., 2004; Rogers, Krebs, Bromley, van der Linden, & Donald, 2006) following a multifractal pattern driven by electrostatic interactions (Vito Foderà, Zaccone, Lattuada, & Donald, 2013). The complexity of this scenario is further enhanced by the fact that *in vitro* amyloid spherulites co-exist with fibrils (V. Foderà & Donald, 2010; Vito Foderà et al., 2013). While the direct observation of fibril growth and time lapses of spherulite growth with temporal resolution of minutes, was recently reported for Aβ peptides (Andersen et al., 2009; Ban et al., 2006; Yagi et al., 2007; Zimmermann et al., 2021), the high heterogeneity of aggregate populations within the same self-assembly reaction presents a further obstacle preventing the correct evaluation of the kinetics of the multiple and concurrent pathways. Indeed, while bulk methods guarantee that the overall propensity of proteins to form amyloid structures is rapidly screened (Vetri & Foderà, 2015), they provide limited information on the aggregation kinetics of individual species, in the form of either fibrils or spherulites, averaging the effect of the morphological heterogeneity of the aggregate population. Consequently, novel methods for real-time and direct observations of single-aggregate growth that can be used to develop and inform models accounting for the heterogeneous growth are highly desirable.

Here we initially combined astigmatism-based 3D direct stochastic optical reconstruction microscopy (dSTORM) (Huang, Wang, Bates, & Zhuang, 2008), spinning disk confocal microscopy (Hayashi & Okada, 2015), and scanning electron microscopy (SEM) to directly observe the formation of individual protein amyloid structures using human insulin (HI) as a model system. Our results allowed us to differentiate among the different species in solution and decipher the nature, morphology and abundance of individual spherulites at different growth stages. Surprisingly, we found that HI spherulite growth is not exclusively isotropic and may occur anisotropically. We developed a method for the detection of Real-time kinetics via binding and Photobleaching Localisation Microscopy (**REPLOM**) to attain real-time videos of the spherulite growth process and reconstruct super-resolution images of the spherulites and their growth kinetics. Using homemade software based on Euclidian minimum spanning tree and machine learning clustering (Jensen et al., 2021; Malle et al., 2021; Pinholt, Bohr, Iversen, Boomsma, & Hatzakis, 2021; Stella et al., 2018; J. Thomsen et al., 2020), we quantitatively associated the growth rates to specific morphological transitions during growth, eventually extracting detailed energy barriers and, thus, the energy landscape for each type of aggregation morphology. We anticipate that the framework presented here will serve as a unique and generic methodology for the simultaneous detection and analysis of multiple species within a single self-assembly reaction. In the specific case of protein systems, the aggregation of which is related to degenerative diseases, our approach provides a platform for connecting kinetics, morphological transitions, and structure and further aid our understanding on interventions against degenerative diseases.

## Results

### Direct observation of diverse structures of HI spherulites by 3D dSTORM, SEM and spinning disk microscopy

We thermally induced insulin amyloid aggregation using an established protocol (Vito Foderà, van de Weert, & Vestergaard, 2010) and examined the bulk kinetics by detecting the fluorescence of the amyloid-sensitive dye Thioflavin T (ThT) and the turbidity signal (Figure S1a). The kinetics traces at incubation temperature of 60 oC, show the classical three-step profile, with the reaction reaching completion after only 3-4 hours. The turbidity and ThT signal perfectly overlapped, suggesting that the aggregation reaction was entirely of an amyloid-like origin (Vito Foderà et al., 2009). Cross-polarized microscopy recordings of the characteristic Maltese cross, indicating spherulite formation under these conditions (Mark R. H. Krebs et al., 2004) (Figure S2). However, standard analysis of the bulk ThT signal was unable to provide information on the morphological transition occurring during the reaction.

To observe directly and with high-resolution the diverse structures of insulin aggregates, we combined the insights obtained from SEM and 3D dSTORM. Using 3D dSTORM allowed us to extend beyond diffraction-limited imaging by TIRF microscopy, which may mask spherulite shape and growth directionality (Huang, Wang, et al., 2008) (Figure 1). Recordings at incubation times between 0.5 to 4 hours points (see methods) provided direct recordings of the diverse early species that can co-exist at the same incubation time. We found spherical-like protein condensates of approximately 200 nm in diameter formed after 0.5 hours, while, a linear pattern was observed with incubation times ranging between 0.5-1h. Surprisingly, the recordings beyond the diffraction limit revealed that at longer incubation times the commonly observed spherulites were found to co-exist in the mixture with anisotropically grown structures (Figure 1, Figure S4, S5c and S5d).

**Figure 1.**
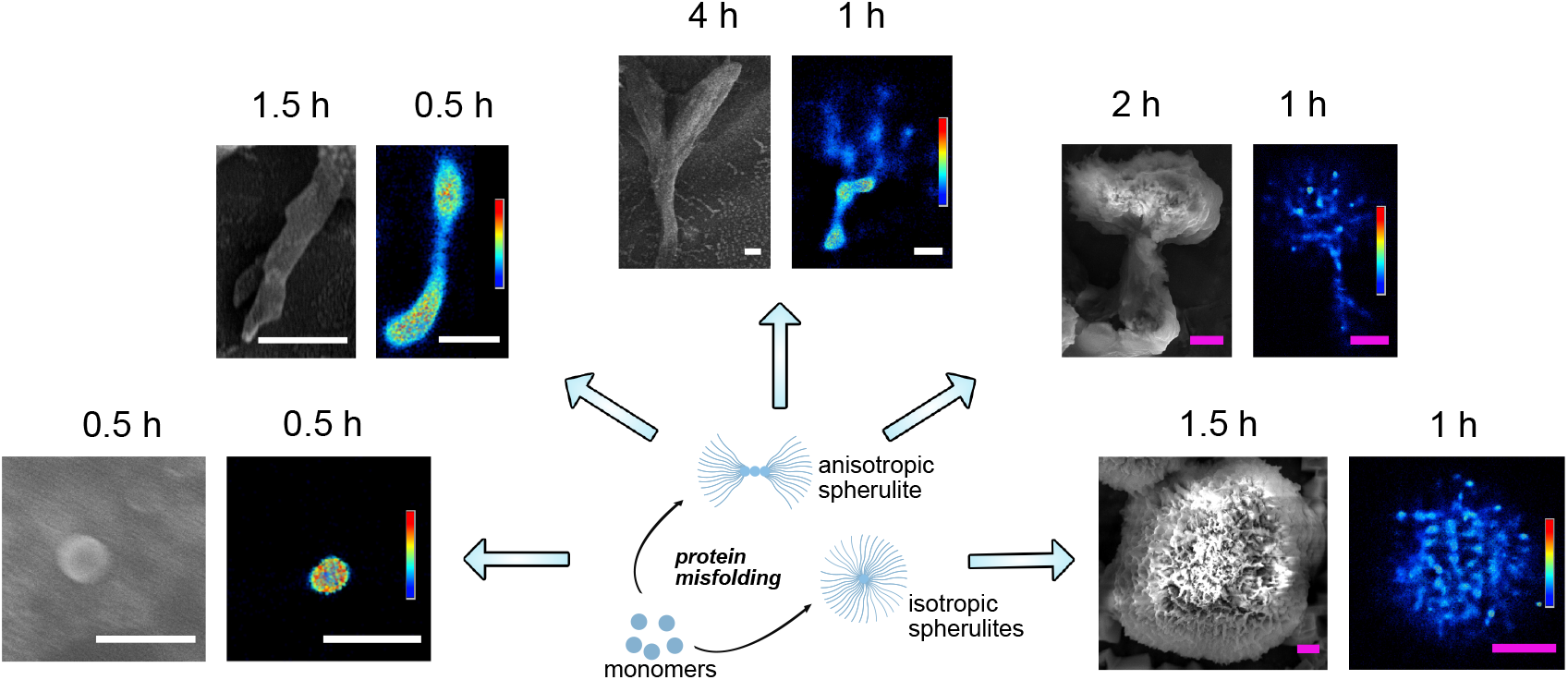
SEM and super-resolution 3D dSTORM reconstructed images of the co-existing in solution morphologies of the anisotropically/isotropically grown structures of different HI aggregates. 3D dSTORM images: density plots, pseudocolor scale corresponds to neighbor localisations: the density of neighboring events within a 100-nm radius sphere from localization. Pseudocolor scale ranges from 0 to 1000 for the first image on left and 0 to 400 for the rest. Scale bars in white color are 1 µm and scale bars in purple color are 5 µm.

The fact that both SEM and 3D dSTORM methods identified the same particle morphology supports this not to be an artifact of fluorescence microscopy, fluorophore labeling (Figure S1b), sample drying for SEM imaging (Figure 1). Note however that depending on conditions the distribution of morphologies may vary slightly consistent with earlier reporting of electrostatic effect for Aβ-(1-40) (Ban et al., 2006) (see Figure S5f). Extending beyond the diffraction limit suggest that protein spherulite growth may diverge from isotropically grown in space (Vito Foderà et al., 2013; Mark R. H. Krebs et al., 2004), and proceed in a preferential direction.

The density plots created with 3D dSTORM (Figure 1) clearly showed that the core had a much higher density than the branching parts, consistent with previous suggestions of the existence of a low-density corona in spherulite structures (Mark R. H. Krebs et al., 2004; Rogers et al., 2006). The formation of the high-density cores appears to indicate the nucleation point, with the subsequent linear-like elongation and branching of slender threadlike fibrils resembling crystalline growth (Gránásy, Pusztai, Tegze, Warren, & Douglas, 2005; Shtukenberg, Punin, Gunn, & Kahr, 2012). This is consistent with the recently proposed initial protein condensation process (Shen et al., 2020), and further growth is determined by tight fibril packing, which forces the biomolecular assembly to occur anisotropically along one specific direction. Delineating this however would require additional experiment and is beyond the scope of this study. The directly observed anisotropy challenges the isotropic spherulite growth, for which the process occurs via the formation of a radiating array of fiber crystallites, but it is observed in the case of crystalline-coil block copolymer spherulites (Song et al.). The origin of such anisotropy might be due to the occurrence of secondary and heterogeneous nucleation at the aggregate surface (Galkin & Vekilov, 2004; Zimmermann et al., 2021), with different binding efficiencies depending on the aggregate areas. While the data in Figure 1 would be consistent with the secondary nucleation, deciphering this with additional data falls beyond the scope of this work.

The diameter of the early linear aggregates increased as a function of time (Table S1). This indicates that the growth was not limited to end-to-end attachment to the linear aggregate, and lateral aggregation also took place. While this is to a certain extent expected (Zimmermann et al., 2021), the super resolution recordings allowed its quantification. The early central linear structures, with diameters of 400 ± 100 nm (see Table S1), successively branched to form radially oriented amyloid fiber-like structures. The further away from the core, the higher the increase in branching frequency, yielding more space-filling patterns. The dimensions of the corona-like structure were ∼2 μm to >20 μm, as shown in Figure 1.

To exclude that diverse morphologies originate from electrostatic interactions with surface immobilization (Ban et al., 2006; Elsharkawy et al., 2018). We used spinning disk microscopy and SEM to detect the morphology of spherulites at different growth stages in solution (see Figure S5). Consistent with the data displayed in Figure 1, we detect both spherulites with asymmetrically grown (Figure 2a and 2b) and symmetrically grown (Figure 2c and 2d) lobes supporting (see 3D videos of Figure 2a and 2c in Supplementary Movies. S1 and S2). We confirmed that the asymmetric growth was not an artifact of substrate depletion, as spherulites with asymmetric lobes had already formed by 2 hours of incubation (Figure 1). This suggests that growth periods of multiple rates occurred within a single sample (Figure S5), which may be masked in bulk kinetics. Moreover, our data indicated the possibility that growth did not occur entirely isotropically from the central core, but rather, there was initially a preferential direction.

**Figure 2.**
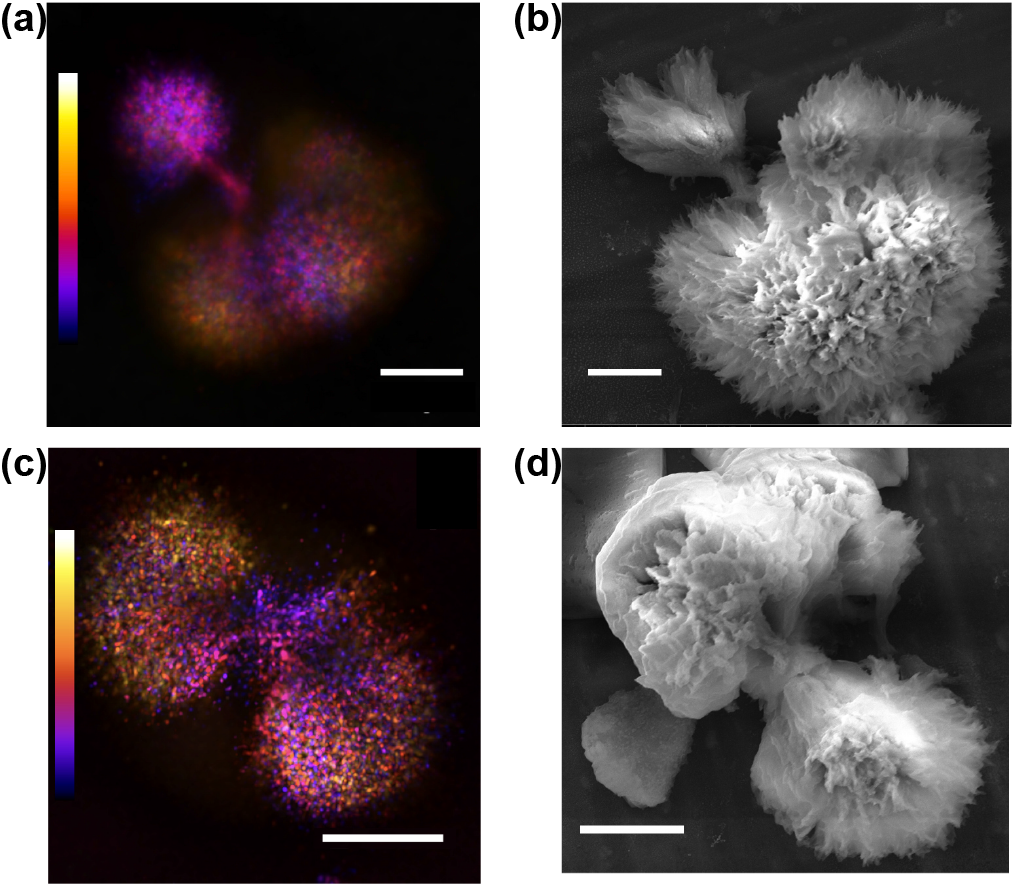
Structure of anisotropically grown human insulin spherulites of two distinct growth-morphologies. a) and b) Spherulites with two asymmetric sides captured by spinning disk confocal microscopy and SEM, respectively. c) and d) Spherulites with symmetric two side structures captured by spinning disk confocal microscopy and SEM, respectively. Data in (a) and (c) were acquired for a sample from incubation time of 16 h at 60 °C. Data in (b) and (d) are for a sample from an incubation time of 4 h at 60 °C. Color scales are from −14.04 μm to 14.04 μm in (a) and −8.46 μm to 8.46 μm in (c). Scale bars are 10 μm. All samples were covalently labeled with Alexa Fluor 647.

### REPLOM: a super resolution method for the real time direct observation of growth of protein aggregation

We developed a new super-resolution experimental method based on single-molecule localisation microscopy, to quantitatively measure the growth rates at the single-aggregate level while simultaneously monitoring the morphological development of the structure. We named the method REal-time kinetics via binding and Photobleaching LOcalisation Microscopy (REPLOM), as it allows researchers, for the first time, to directly image the morphological development of each individual aggregates in real-time with super-resolution and, simultaneously, access the kinetic traces for thermodynamic analysis of the process. To perform REPLOM, HI monomers were covalently labeled with Alexa Fluor 647 NHS Ester (see Supplementary Information for experimental details). Figure 3a illustrates how REPLOM works: initially, only small protein condensates, i.e., cores, are formed and bind to the poly-L-lysine-covered surface. The spatial location of each of the fluorophores is accurately detected prior to their photobleaching (Bohr et al., 2019; Moses et al., 2021). Optimizing the imaging settings and the absence of imaging buffer ensures rapid chromophore bleaching after binding (see Methods and Figure S8). As the growth progresses, additional HI monomers from the solution bind to the core, extending the dimensions of the aggregate. Each labeled insulin binding event results in a diffraction-limited spot, the precise location of which can be accurately extracted, similarly to in photoactivated localization microscopy (PALM) methodologies (Betzig et al., 2006) (see Methods and Figure S9 for resolution of the method and Supplementary Movie S3-S4).

**Figure 3.**
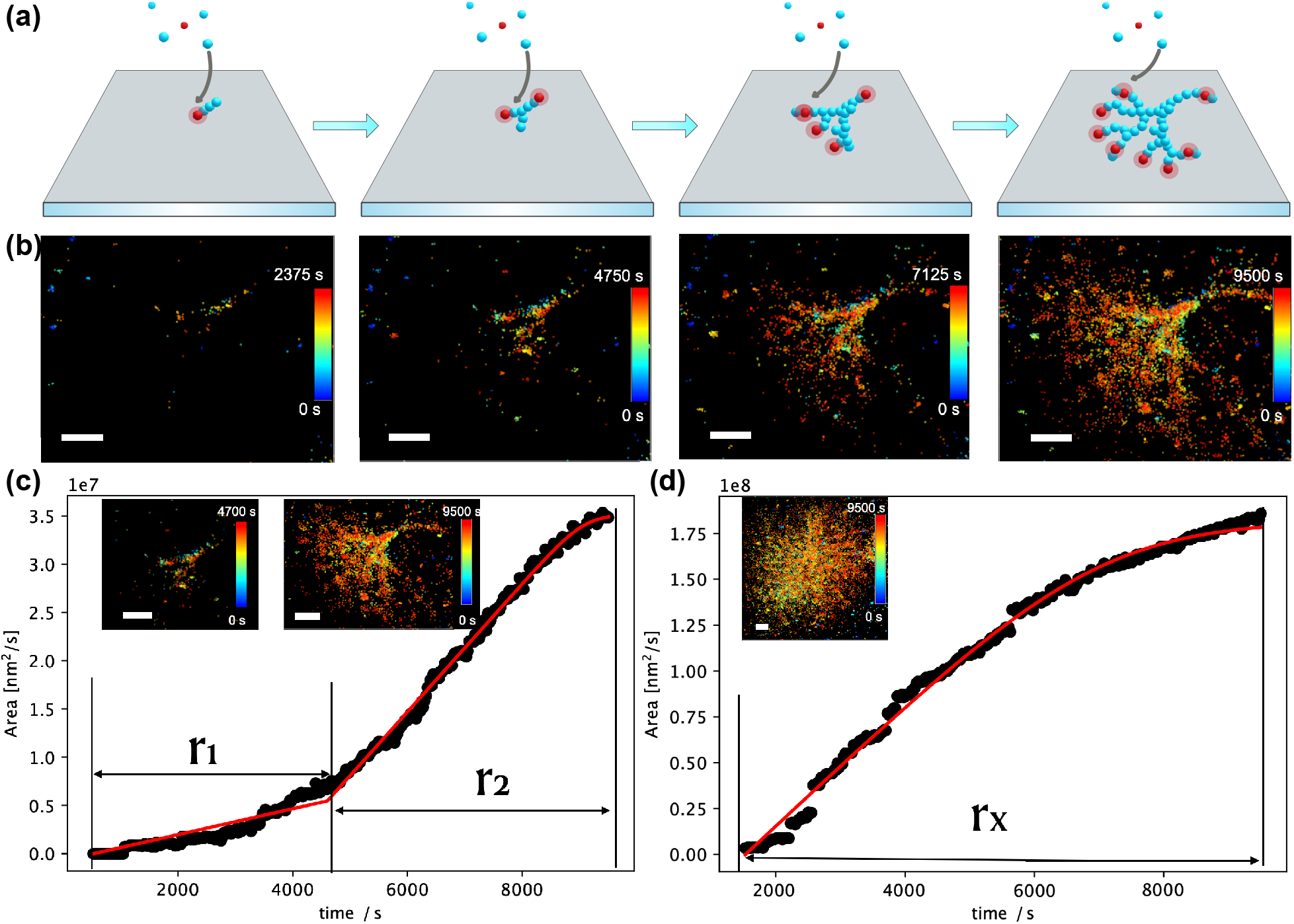
Direct real-time observation of HI aggregate growth by REPLOM (real-time kinetics via binding and photobleaching localisation microscopy). a) Cartoon representation of REPLOM: initially, the fluorescent signal from the small fluorescently labeled protein condensates was detected, followed by chromophore photobleaching. As the growth progressed, labeled insulins from solution bound to the aggregate, increasing the dimensions. Each binding event resulted in a diffraction-limited spot, the coordinates of which were accurately extracted, before it was photobleached by the intense laser. Parallelized recordings of the spatially distinct binding of multiple individual emitters revealed the temporal morphological development of several aggregates (Red is Alexa Fluor 647-labeled HI in fluorescent state, and blue is un-labeled insulin or Alexa Fluor 647-labeled HI in dark/photobleached state). b) Direct real-time observation of temporal development of anisotropic growth at t = 2375 time intervals. Scale bars: 2 μm. c) and d) Growth curves of anisotropic spherulite (c) and isotropic spherulite (d). For anisotropic spherulites, the curve contains two parts roughly correlating with the formation of the core/linear part and branching part (see method REPLOM section). Isotropic spherulite growth was linear and followed by saturation. Inset: the corresponding HI spherulite obtained by REPLOM. Scale bars: 2 μm. See SI for the movies.

Parallelized recordings of the spatially distinct binding of multiple individual HI loaded with emitters allow the real-time direct observation of the temporal morphological development of each aggregate (see Figure 3b, Figure S10, and Supplementary Movies, S3-S4). The methodology is reliant on the intrinsic bleaching of chromophores to extract their coordinates (Burnette, Sengupta, Dai, Lippincott-Schwartz, & Kachar, 2011; Gordon, Ha, & Selvin, 2004; Qu, Wu, Mets, & Scherer, 2004) and is similar to Binding Activation Localisation Microscopy (BALM) (Ries et al., 2013), which measures existing structures, but additionally facilitates real-time direct observation of the growth process. It also extends beyond recent methods based on conventional TIRF to observe exclusively fibril growth (Zimmermann et al., 2021) or low temporal resolution time lapses of linear or spherulite growth (Andersen et al., 2009; Ban et al., 2006; Yagi et al., 2007), offering in addition rate recording and morphological development of both fibrillar and spherulite structures even below the diffraction limit. Consequently, the geometry and morphological development of each aggregate can be observed directly with sub-diffraction resolution, offering the extraction of each particle’s growth kinetics.

### Extraction of growth rates for diverse aggregate morphologies

Consistent with the 3D dSTORM data, the direct observation of HI spherulite growth by REPLOM confirmed that HI spherulites grow both anisotropically and isotropically (Figure 3). To extract the growth rate kinetics for each individual aggregate, we identified the points belonging to the growing aggregate with an approximate Euclidean Minimum Spanning tree segmentation (Cowan & Ivezić, 2008) and estimated the area using a Gaussian mixture model based on hierarchical clustering in Figure 3c and 3d (see Supporting Information for the details) (Jensen et al., 2021; Pinholt et al., 2021; Stella et al., 2018; J. Thomsen et al., 2020). For isotropic morphologies, a single linear growth rate was observed (r_x_) followed by a plateau (see Supporting Information), while for anisotropic morphologies the growth curve consisted of two rate components (r_1_ and r_2_), as shown in Figure 3c and 3d and Figure S11; r_1_ corresponds to the initial linear core and r_2_ to the branching part, and they best fitted to reaction-limited linear growth and a diffusion-limited sigmoidal growth, respectively (Domike & Donald, 2007, 2009; Goldenfeld, 1987; Majumder, Busch, Poudel, Mecking, & Reiter, 2018; Tanaka & Nishi, 1985) (see Methods and Supplementary Movies S3-S8). Consequently, the growth rates (r_1_, r_2_, and r_x_) for each individual aggregate were extracted. The growth readouts of the individual geometrically distinct morphologies allowed us to go beyond the standard analysis of sigmoidal curves, which does not yield information on, or discriminate between, the temporal developments for each morphology. REPLOM revealed that the anisotropic growth operated via a two-step process imposed by the geometry of the growth—a pattern masked in current super-resolution and bulk readouts.

### Extraction of Energy barriers for the growth of diverse HI spherulites morphologies

The real-time single-particle readout from REPLOM facilitates the kinetic analysis of the temperature dependence of growth for each diffraction limited type of spherulite morphology and, consequently, the extraction of the activation energy barriers for both the spherulite morphologies and growth phase. Therefore, HI aggregate formation was induced at three different temperatures accessible without introducing optical artefacts in our microscopy setup: 45 oC, 37 oC, and 32 oC. The ThT fluorescence measurements at the three temperatures representing the average growth kinetics are shown in Figure 4a. The rate distributions at the three temperatures for each type of morphological growth are shown in Figure 4 b, 4c, and 4d (N = ∼20, see also Figure S12). As expected, the linear parts r_1_ (Figure 4b) and branched parts r_2_ (Figure 4c), as well as the isotropic growth rate r_x_ (Figure 4d), increased at increased incubation temperature. The data do not show a pronounced curvature, and this may be due to the narrow temperature range investigated in our study and is in agreement with earlier studies(Buell et al., 2012). This would suggest that the differences in heat capacity between the soluble states of the proteins and the transition states for aggregation are small (Buell et al., 2012). Using the Arrhenius equation (Buell et al., 2012; Cohen et al., 2018) (Figure 4e-4h), we extracted the activation energy of each of the isotropic or anisotropic morphological growths and the respective linear or branching part of the individual aggregates. For the linear part of the anisotropic spherulites, the activation energy was 104.67 ± 10.23 kJ/mol (Figure 4f), while for the branched part it was 84.05 ± 5.46 kJ/mol (Figure 4g), and for the isotropically grown spherulites it was 87.25 ± 6.44 kJ/mol (Figure 4h). The activation energy extracted from the bulk kinetics shown in Figure 4e (93.95 ± 14.91 kJ/mol) is consistent with data on bovine insulin fibril formation (∼100 kJ/mol) (Buell et al., 2012). The REPLOM methodology on the other hand allowed deconvolution of a higher barrier related to step 1 in the anisotropic growth (r_1_) and lower barrier in the branching part of isotropic and anisotropic growth (r_x_ and r_2_). Together, these data indicate that the pronounced heterogeneity of growth mechanisms and structures within the aggregation ensemble leads to heterogeneity of the activation barriers. We indeed highlighted that spherulite growth may proceed both isotropically and anisotropically, with the latter presenting a two-step process imposed by the geometry of the growth and characterized by two activation energies that are markedly different to those obtained by bulk kinetics and for insulin fibrils (Buell et al., 2012).

**Figure 4.**
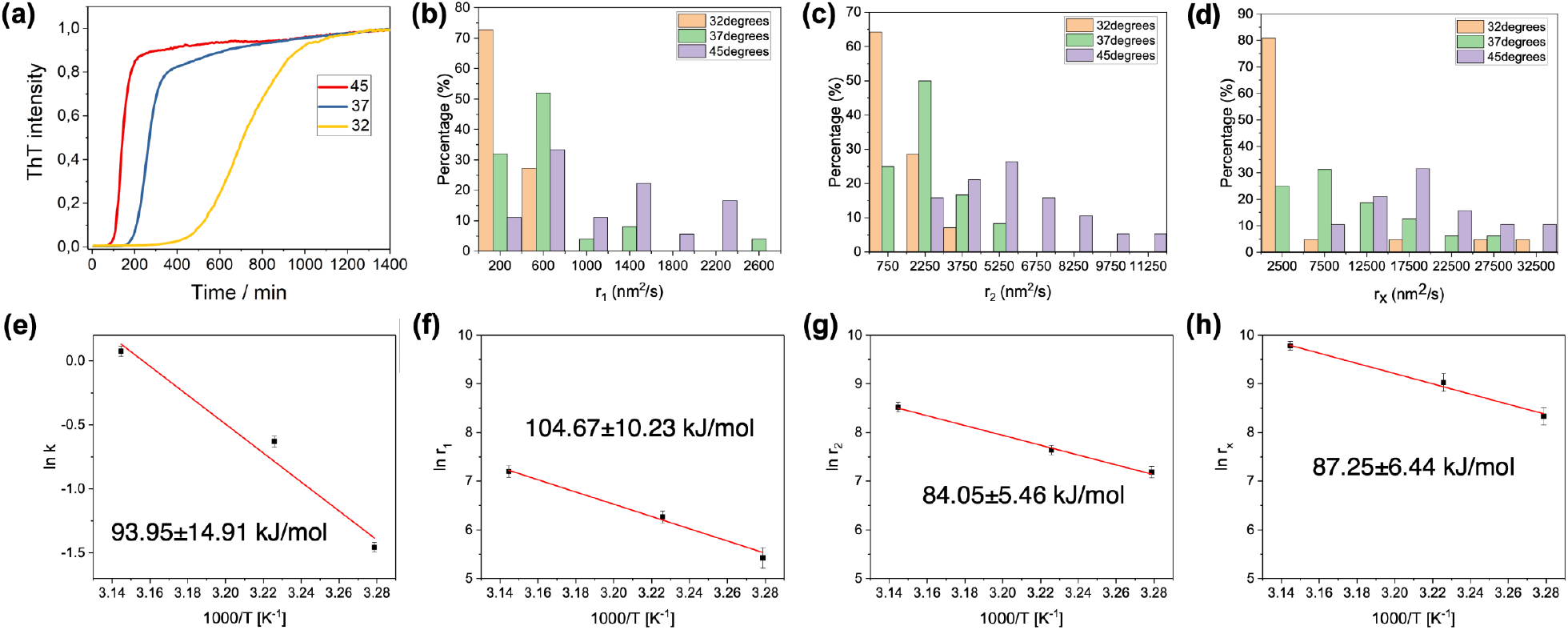
Kinetic and thermodynamic characterization of insulin aggregation. a) Normalized bulk ThT fluorescence kinetics on with incubation temperatures of 45, 37, and 32 °C. b) and c) REPLOM-extracted rate distribution of anisotropic aggregates at the three different incubation temperatures: (b) linear part and (c) branching part. d) Rate distribution of isotropic aggregates at the three different incubation temperatures. e-h) Arrenhius plots for spherulites obtained from bulk experiments (e), and REPLOM (f-h). The formation of linear (f) and branched (g) parts of anisotropic spherulites, and the formation of isotropic spherulites (h).

## Discussions

Our combined results revealed that the growth of amyloid core-shell structures for insulin, i.e., spherulites, may proceed not only via isotropic growth but also by following a multistep pathway characterized by initial pronounced anisotropic behavior (Figure 5). The anisotropic growth may thus not be an exclusive property of metal alloys, salts and minerals, but may extend to protein aggregates. In essence are data are consistent with a unifying mechanism underlying chemical growth of both biological soft materials and hard-non biological composites. Such variability in growth within the same aggregation reaction results in a spectrum of aggregation kinetics traces that can be quantitatively detected by our method, allowing the operator to extract the thermodynamic parameters for each of the aggregation subsets. These findings underscore how conclusions solely based on bulk kinetics data may overlook the complexity and heterogeneity of the aggregation process.

**Figure 5.**
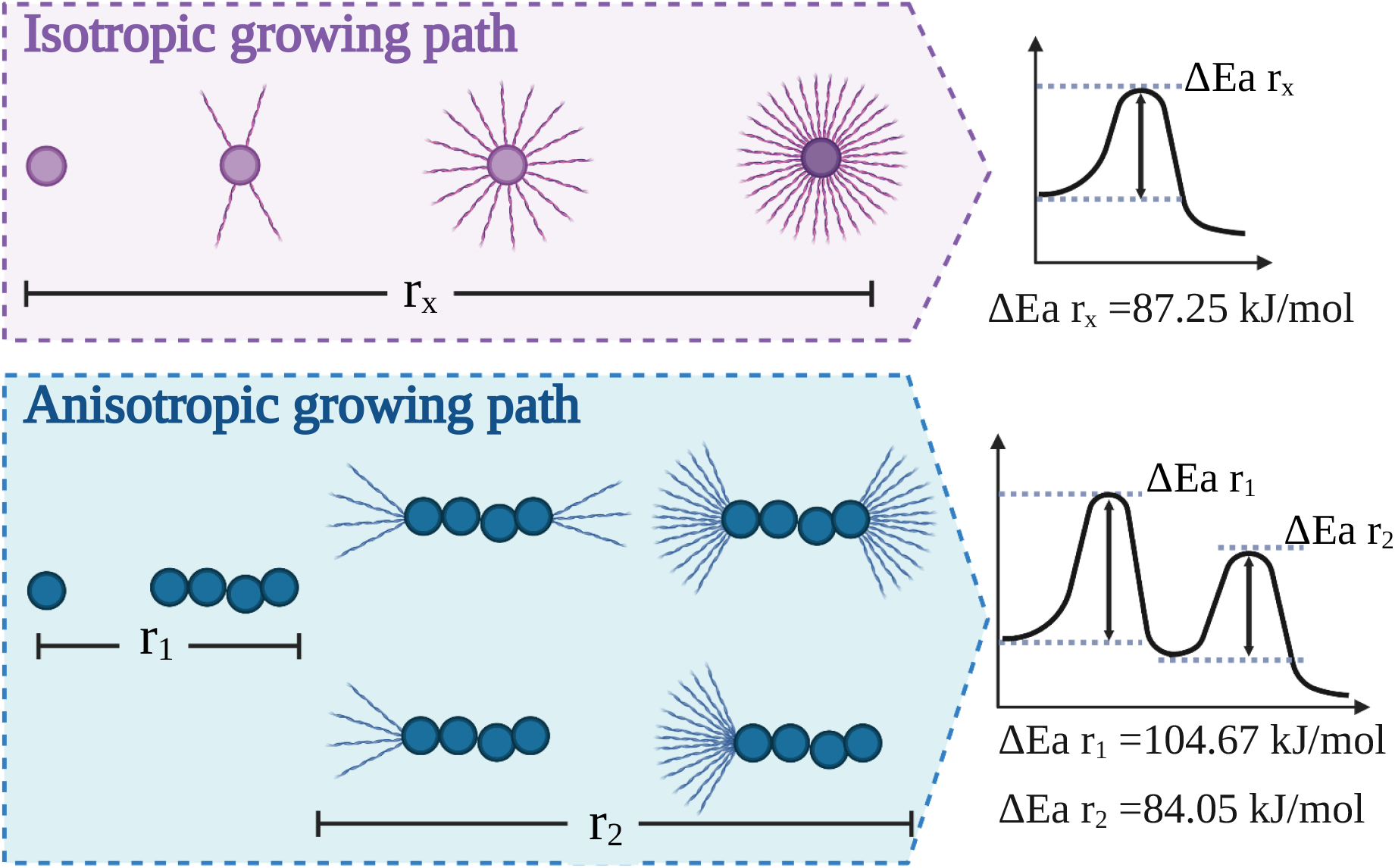
Schematic representation of the diverse pathways of insulin aggregation and their respective energy barriers. Top: Isotropic spherulite growth, where fibril-like filaments isotropically and radially grow on a dense core. Process is characterized by a single activation energy of ∼87 kJ/mol. Bottom: anisotropic growth, where the dense core is growing linearly before it successively branches to form radially oriented amyloid fiber-like structures. The further the branching from the core, the more increased the branching frequency, yielding a more space-filling pattern. The process involves two steps imposed by the geometry of the growth and characterized by two activation energies of 104 and 84 kJ/mol for the linear and branching parts, respectively.

Our novel experimental approach offers real-time detection of super-resolution images during protein aggregation kinetics. The REPLOM method allows the direct observation of self-assembly kinetics at the level of single aggregates and the quantification of the heterogeneity of aggregates and their growth mechanisms, which are otherwise masked with current methodologies. Our general framework can be extended to the simultaneous detection of markedly different structures within a single aggregation reaction and contribute to research into a more comprehensive representation of the generalized energy landscape of proteins. This will offer the unique possibility of disentangling different mechanisms leading to the myriad of aggregate structures that occur. The method is implemented on the insulin model systems, but can be easily translatable to more medically relevant proteins, such as α-synuclein or Aβ peptide. Deciphering whether these structures persist in the context of the cellular environment and the direct physiological implications of anisotropically grown morphologies would require combination of our methodologies with DNA-paint and antibodies as recently developed (Sang et al., 2021). Our approach may indeed provide unprecedented information on transient intermediate species, which are nowadays recognized as the cause of progression in many diseases, in terms of both energetics and morphology. Finally, our approach is general and may be applicable to generic self-assembly reactions of systems characterized by a high degree of heterogeneity.

## Materials and Methods

### Human insulin (HI) labeling and spherulite preparation

Alexa Fluor 647 NHS Ester (ThermoFisher Scientific) was dissolved in anhydrous-DMSO to a concentration of 2 mg/mL. 5 μL of the dye solution was added to 1 mL 5 mg/mL HI (91077C, Sigma-Aldrich, 95%) monomer solution, mixed gently and thoroughly. The mixed solution was allowed to react for ∼2 hours at room temperature to complete the conjugation. After that the labeled protein was purified from the excess of free dye by a PD SpinTrap G-25 column (GE Healthcare), divided into aliquots and stored at -80 °C.

HI spherulites were formed in 0.5 M NaCl, 20% acetic acid (VWR Chemicals, 98%) solution with pH around 1.7. The ratio of labeled to unlabelled HI monomer was about 1 to 60000 (dSTORM) or 1 to 10000 (RE-PLOM), with the final concentration of HI was 5 mg/mL. The solution was filtered through 0.22 μm filters (LABSOLUTE) and then incubated in a block heater.

### Atto 655-labeled liposome preparation for 3D dSTORM calibration

Atto 655-labeled liposomes with 2% negative charge which were used for the 3D dSTORM calibration were prepared as previously published method (R. P. Thomsen et al., 2019). In detail, a ratio of 97/2/0.5/0.5 for 1,2-Dioleoyl-sn-Glycero-3-Phosphocholine (DOPC), 1,2-dioleoyl-sn-glycero-3-phospho-L-serine (sodium salt) (DOPS), 1,2-distearoyl-snglycero-3-phosphoethanolamine-N-[biotinyl(polyethylene glycol)-2000] (ammonium salt) DSPE-PEG_2000_-biotin and Atto655-PE were added to a glass vial. The solvent chloroform was removed completely by nitrogen flow for about 10 minutes followed by vacuum for several hours. The lipid film was rehydrated in MES buffer (pH 5.6) to final total lipid concentration 0.5 mg/mL, vortexed for 30 seconds and incubated for 30 minutes. The sample was extruded 11 times through a polycarbonate membrane filter with a pore size 50 nm. Then the liposome suspension was exposed to 10 cycle of flash-freezing and thawing so as to ensure an unilamellar membrane structure. The liposomes were aliquoted and stored at -20 °C.

### Turbidity / Thioflavin T (ThT) fluorescence kinetics

For in situ absorbance or ThT fluorescence, experiments were carried out using a plate reader system (BMG LABTECH, CLARIOstar) with 96-microwell polystyrene plates (Nalge Nunc, ThermoFisher Scientific). Each well contained 200 μL solution. The plates were covered with a self-adhesive sealing film (nerbe plus, for absorbance) or a clear polyolefin film with sealing tape (Thermo Fisher Scientific, for ThT fluorescence) to avoid evaporation of the samples and incubated at the desired temperatures without mechanical shaking. For absorbance, the excitation wavelength was 480 nm; and for ThT fluorescence, the solution contained 20 μM ThT and the emission intensity at 486 nm was recorded upon excitation at 450 nm. The signal was detected every 309 s.

### Spinning Disk Microscopy

The 3D images of grown spherulites were taken by a SpinSR10-spinning disk confocal super resolution microscope (Olympus) using a silicone oil-immersion 100x objective (UPLSAPO100XS, NA=1.35, Olympus). The Alexa Fluor 647-labeled HI spherulites were excited with a 640 nm laser (OBIS COHERENT). The exposure time was 50 ms and the z step length was 0.36 μm.

### Scanning Electron Microscopy (SEM)

SEM images of spherulites were taken by using a Quanta FEG 200 ESEM microscope.

### Cross Polarized Microscopy

Images were collected using a 10x objective and crossed polarised which enabled spherulites to show the characteristic Maltese cross (Zeiss Axioplan Optical Microscope, Carl Zeiss).

### Super-resolution Imaging

Super resolution imaging was attained on an inverted Total Internal Reflecion microscope (TIRF) (Olympus IX-83) with a 100x oil immersion objective (UAPON 100XOTIRF, NA=1.49, Olympus) Alexa Fluor 647 was excited by a 640 nm solid state laser line (Olympus) and reflected to a quad band filter cube (dichroic mirrors ZT640rdc, ZT488rdc and ZT532rdc for splitting and with single-band bandpass filters FF02-482/18-25, FF01-532/3-25 and FF01-640/14-25). Signal was detected by an EMCCD camera (imagEM X2, Hamamatsu).

### 3D direct Stochastic Optical Reconstruction Microscopy (3D dSTORM) and image analysis

3D dSTORM imaging was achieved by installing a cylindrical lens (f = 500 mm) in the emission pathway of (TIRF) to introduce the astigmatism of point spread function (PSF) (Huang, Wang, et al., 2008). All the dSTORM imaging experiments were performed at room temperature (21 °C). The exposure time was 30 ms and 10000 frames for each movie.

To extract z information from the widths of single molecule images, we generated a calibration curve of PSF width in the lateral plane (Wx and Wy) as a function of height by measuring Atto 655-labeled liposomes using TIRF with a step size of 10 nm and exposure time of 30 ms (Fig. S3).

The HI aggregates which were incubated in a block heater for 0.5 hours to 2 hours at 60 °C. At the desired time they were added to the poly-L-Lysine treated microscope chamber (Chen et al., 2017) and incubated for 10 min at room temperature to ensure immobilization. Extra sample was washed away with MilliQ water. Imaging buffer containing 50 mM Tris, 10 mM NaCl, 10% (w/v) glucose, 0.5 mg/mL glucose oxidase, 40 μg/mL catalase and 0.1 M MEA (Huang, Jones, Brandenburg, & Zhuang, 2008) was flushed into the chamber for dSTORM imaging. All measurements were carried out at room temperature. The optimal ratio of labeled to unlabeled insulin that provided reliable signal without affecting the aggregation process or compromising resolution was 1 to 60,000. This is quite different from earlier dSTORM imaging of fibrils using a ratio of 1/20 (Pinotsi et al., 2014) because of the much higher 3D density of spherulites that prevent reliable super resolution imaging at high labeling ratios.

The 3D dSTORM data was analysed by ThunderSTORM (Ovesný, Křížek, Borkovec, Švindrych, & Hagen, 2014). The z information of individual localisations was extracted based on the calibration curves (calculated by ThunderSTORM, shown in Figure S3). The detected localisations were further filtered according to their intensity and drift correction, in order to remove some possible false positive or poor quality detections. 3D super-resolution images were visualized with ViSP software (Beheiry & Dahan, 2013).

### REal-time kinetic via Photobleaching Localisation Microscopy (REPLOM)

#### Preparation of HL aggregates and imaging

The solution containing 5 mg/mL HI monomer was first incubated in a block heater to skip the lag phase. The optimal pre-incubation time for spherulite formation on the microscope surface was found to be ∼ 8 hours for 45 oC, 20 hours for 37 oC and 75 hours for 32 oC, respectively. Then they were transferred to poly-L-lysine coated glass slide chambers and covered by a lip to prevent solvent evaporation during imaging (Figure S6).

REPLOM was performed on the same TIRF microscope setup as the 3D STORM without the cylindrical lens. Alexa Fluor 647 labeled HI was excited by 640nm solid state laser lines (Olympus). We found the optimal ratio of labeled to unlabeled insulin for REPLOM to be ca. 1 to 10,000. A high labelling density would result in proximate fluorophores from the newly grown area emitting simultaneously and therefore cause mislocalization (Pinotsi et al., 2014). Too low labeling ratio may cause some details, e.g. small branching part, during spherulites growth to be undetected. Imaging was performed with an exposure time of 30 ms followed by a waiting time for each frame of 20-40 seconds so as to capture in real time the slow kinetics of spherulite formation. This frame rate allowed to capture both seed formation and extract the growth rate of insulin aggregates. Faster frame rates may be required for different protein aggregates (Ogi et al., 2014). All image acquisition was performed at the same incubation temperatures as in the block heater. The incubation temperatures during the imaging processes were achieved by a heating unit 2000 (PECON).

### Data analysis

The data was analysed by ThunderSTORM. Some possible false positive or poor-quality detections were removed by intensity filter. Figure S7 shows the comparison of images prior to and after drift correction. The reconstructed images with time series were obtained by ViSP (Beheiry & Dahan, 2013) software. For Quantification of growth kinetics is available in Supporting Information.

### Lifetime of fluorophores

The lifetime of fluorophores in REPLOM was evaluated by checking the duration time of fluorescent state before they were photobleached. We checked 1885 individual Alexa Fluor 647 fluorophores and found they were photobleached very fast without imaging buffer (Figure S7). The lifetime is about 0.7845 ± 0.0017 frames.

### Resolution of REPLOM

The resolution of REPLOM was determined by the FWHM of single spot’s intensity (Figure S9) using an adapted version of previously published software (Bohr et al., 2019; R. P. Thomsen et al., 2019). Briefly, using our subpixel resolution software, we were able to extract multiple (91) single spots (see Figure S9) and align all to the same center. Fitting a two-dimensional gaussian to the resulting stacked clusters allowed the reliable extraction of FWHM used to determine the obtained resolution. Using a maximum likelihood fitting scheme avoided potential bias from data binning.

### Quantification of growth kinetics by Euclidean Minimum Spanning tree

The method for identification of candidates for fluorophores docking on a growing aggregate was inspired by recent published work (Cowan & Ivezić, 2008) and done in the following way:

First, using all detected RE-PLOM spots from the movie, an approximate Euclidean Minimum Spanning tree was constructed using only the 30 nearest neighbors as candidates for edges. Regions of aggregate candidates were cut from each other by removing all edges with lengths more than the 95th percentile. This is an effective way of separating high-density regions from low-density regions. The computation was done using the function HierarchicalClustering from the astroML python package. Since we were interested mostly in the large insulin aggregates where the internal structure was visible, it was decided that all clusters obtained in this manner with less than 100 detected fluorophores were excluded from the subsequent analysis.

The time-dependency of the aggregate growth was found by a similar approach. At each frame, for a cluster, a refined grouping was done by cutting an approximate Euclidean Minimum Spanning tree made using 10 neighbors with a distance cutoff of 400nm which was found to be optimal for removal of most spots outside the aggregate while still not cutting up the main group. The points from the largest subgroup resulting from this analysis were defined to be the aggregate for that frame.

The area of the aggregate was estimated using a gaussian mixture model with a component for every 5 points in the aggregate, but not less than 25 components (Cowan & Ivezić, 2008). We defined the area of the aggregate as the region lying above the average probability density in this fit. The growth profile resulting from our approach had a few artifacts like jumps and fluctuations due to mixture model fitting and aggregate segmentation, but we found that the resulting growth curve in most cases had an identifiable trend, and the results were quite consistent across parameter choices.

From the estimated area of the aggregate in each frame, a growth curve could be plotted.

The radial growth rate of such aggregates has previously been found to be either reaction-limited or diffusion-limited, leading to linear increase in time or increase as 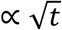 respectively (Domike & Donald, 2007; Goldenfeld, 1987; Majumder et al., 2018; Tanaka & Nishi, 1985). If we assume that the estimated area of the aggregated is directly related to the radius as *A* ∝ *R*^2^ the two growth types lead to the following models

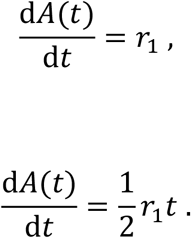

Where the first model is diffusion limited and the second is reaction limited. We found that many of the structures where initially consistent with reaction limited diffusion and then shifted to either diffusion or reaction limited growth with a new rate. To allow for this shift, we let the growth be diffusion limited up to a switch-point *t*_2_ after which the growth rate changes. We formulate one such model which ends reaction limited and one which remains diffusion limited

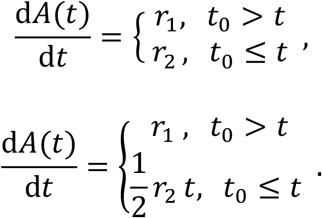

Finally, without continuous flow of constituent monomer, the growth inevitably saturates at a plateau (Domike & Donald, 2009). For both models, we therefore introduce a switch time *t*_1_ after which the growth slowly saturates sigmoidally over a time interval 5τ

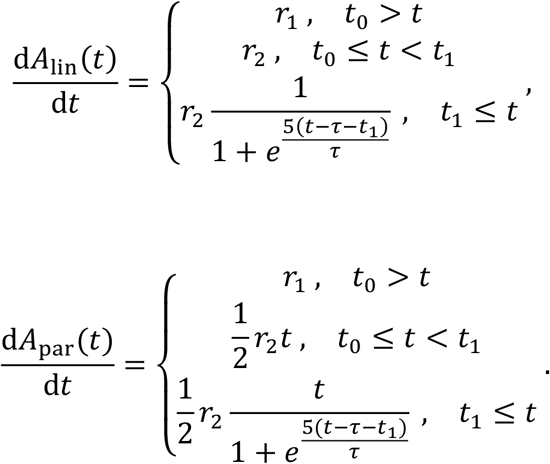

Where we introduced the names *A*_lin_ and *A*_par_ referring to the linear-like and parabolic-like shape of the two resulting growth curves. We found the anisotropic spherulites to fit best with *A*_lin_ and the isotropic spherulites fit best with *A*_par_.

When fitting an experimentally observed aggregate growth curve {*A*_*i*_, *t*_*i*_}, *i* ∈ (0, *N* − 1) the equations where numerically integrated from an initial timepoint (*A*_0_, *t*_0_)to the final timepoint (*A*_*N* − 1_, *tN* − 1). For each growth curve, the parameters (*r*_1_, *r*_2_, *t*_0_, *t*_1_, τ)were estimated with a chi2 fit. Each fit was run twice, the first fit was unweighted and were used to estimate the error bars using the standard deviation of the residuals. The second fit used the residuals in a weighted chi2 fit to obtain the final fit parameters for the growth curve.

## Data availability

All data sets used for figures are provided as source data in the manuscript. Source code and executable can be found at https://github.com/hatzakislab/REPLOM-analysis-tool. All source data are available at https://sid.erda.dk/sharelink/fje3exOlq2.

## Author information

### Author Contributions

M.Z, N.S.H and V.F wrote the paper with feedback from all authors. M.Z designed, carried out and analysed all microscopy experiments, and prepared all samples. H.D.P wrote the automated cluster finding and rates analysis algorithm. M.Z and X.Z did the ThT-fluorescence and turbidity measurements. S.S-R.B calculated the resolution of REPLOM and fluorophore’s lifetime. L.B and A.Z helped with the mechanism explanation. N.S.H conceived the project idea, in collaboration with V.F., and had the overall project management and strategy.

## Notes

The authors declare no competing financial interest.

## Acknowledgment

This work was funded by the Lundbeck foundation (grant R250-2017-1293 and R346-2020-1759) for M.Z. Villum foundation young investigator fellowship (grant 10099), and the Carlsberg foundation Distinguished Associate professor program (CF16-0797) and the NovoNordisk Center for Biopharmaceuticals and Biobarriers in Drug Delivery (NNF16OC0021948) for N.S.H. Villum foundation young investigator fellowship (grant 19175), the Novo Nordisk foundation (NNF16OC0021948) and Lundbeck foundation (R155-2013-14113) for V.F. China Scholarship Council (201709110108) for X.Z. Work at The Novo Nordisk Foundation Center for Protein Research (CPR) that NSH is associated with, is funded by a generous donation from the Novo Nordisk Foundation (Grant number NNF14CC0001). We thank Dr Y. Hu from Technical University of Denmark for the help with SEM imaging.

N.S.H. and V.F are members of the Integrative Structural Biology Cluster (ISBUC) at the University of Copenhagen.

## REFERENCES

Andersen, C. B., Yagi, H., Manno, M., Martorana, V., Ban, T., Christiansen, G., … Rischel, C. (2009). Branching in Amyloid Fibril Growth. Biophysical Journal, 96(4), 1529–1536. doi:http://doi.org/10.1016/j.bpj.2008.11.024

Ban, T., Morigaki, K., Yagi, H., Kawasaki, T., Kobayashi, A., Yuba, S., … Goto, Y. (2006). Real-time and Single Fibril Observation of the Formation of Amyloid β Spherulitic Structures. Journal of Biological Chemistry, 281(44), 33677–33683. doi:10.1074/jbc.M606072200

Beheiry, M. E., & Dahan, M. (2013). ViSP: representing single-particle localizations in three dimensions. Nature Methods, 10, 689. doi:10.1038/nmeth.2566 https://www.nature.com/articles/nmeth.2566#supplementary-information

Betzig, E., Patterson, G. H., Sougrat, R., Lindwasser, O. W., Olenych, S., Bonifacino, J. S., … Hess, H. F. (2006). Imaging Intracellular Fluorescent Proteins at Nanometer Resolution. Science, 313(5793), 1642–1645. doi:10.1126/science.1127344

Bohr, S. S. R., Lund, P. M., Kallenbach, A. S., Pinholt, H., Thomsen, J., Iversen, L., … Hatzakis, N. S. (2019). Direct observation of Thermomyces lanuginosus lipase diffusional states by Single Particle Tracking and their remodeling by mutations and inhibition. Scientific Reports, 9(1), 16169. doi:10.1038/s41598-019-52539-1

Buell, A. K., Dhulesia, A., White, D. A., Knowles, T. P. J., Dobson, C. M., & Welland, M. E. (2012). Detailed Analysis of the Energy Barriers for Amyloid Fibril Growth. Angewandte Chemie International Edition, 51(21), 5247–5251. doi:https://doi.org/10.1002/anie.201108040

Burnette, D. T., Sengupta, P., Dai, Y., Lippincott-Schwartz, J., & Kachar, B. (2011). Bleaching/blinking assisted localization microscopy for superresolution imaging using standard fluorescent molecules. Proceedings of the National Academy of Sciences, 108(52), 21081–21086. doi:10.1073/pnas.1117430109

Chen, W., Young, L. J., Lu, M., Zaccone, A., Ströhl, F., Yu, N., … Kaminski, C. F. (2017). Fluorescence Self-Quenching from Reporter Dyes Informs on the Structural Properties of Amyloid Clusters Formed in Vitro and in Cells. Nano Letters, 17(1), 143–149. doi:10.1021/acs.nanolett.6b03686

Chiti, F., & Dobson, C. M. (2006). Protein Misfolding, Functional Amyloid, and Human Disease. Annual Review of Biochemistry, 75(1), 333–366. doi:10.1146/annurev.biochem.75.101304.123901

Cohen, S. I. A., Cukalevski, R., Michaels, T. C. T., Šarić, A., Törnquist, M., Vendruscolo, M., … Linse, S. (2018). Distinct thermodynamic signatures of oligomer generation in the aggregation of the amyloid-β peptide. Nature Chemistry, 10(5), 523–531. doi:10.1038/s41557-018-0023-x

Cowan, N. B., & Ivezić, Ž. (2008). The Environment of Galaxies at Low Redshift. The Astrophysical Journal, 674(1), L13–L16. doi:10.1086/528986

Domike, K. R., & Donald, A. M. (2007). Thermal Dependence of Thermally Induced Protein Spherulite Formation and Growth: Kinetics of β-lactoglobulin and Insulin. Biomacromolecules, 8(12), 3930–3937. doi:10.1021/bm7009224

Domike, K. R., & Donald, A. M. (2009). Kinetics of spherulite formation and growth: Salt and protein concentration dependence on proteins β-lactoglobulin and insulin. International Journal of Biological Macromolecules, 44(4), 301–310. doi:https://doi.org/10.1016/j.ijbiomac.2008.12.014

Elsharkawy, S., Al-Jawad, M., Pantano, M. F., Tejeda-Montes, E., Mehta, K., Jamal, H., … Mata, A. (2018). Protein disorder-order interplay to guide the growth of hierarchical mineralized structures. Nature Communications, x9. doi:ARTN 214510.1038/s41467-018-04319-0

Exley, C., House, E., Collingwood, J. F., Davidson, M. R., Cannon, D., & Donald, A. M. (2010). Spherulites of Amyloid-beta(42) In Vitro and in Alzheimer’s Disease. Journal of Alzheimers Disease, 20(4), 1159–1165. doi:10.3233/jad-2010-091630

Foderà, V., Cataldo, S., Librizzi, F., Pignataro, B., Spiccia, P., & Leone, M. (2009). Self-Organization Pathways and Spatial Heterogeneity in Insulin Amyloid Fibril Formation. The Journal of Physical Chemistry B, 113(31), 10830–10837. doi:10.1021/jp810972y

Foderà, V., & Donald, A. M. (2010). Tracking the heterogeneous distribution of amyloid spherulites and their population balance with free fibrils. The European Physical Journal E, 33(4), 273–282. doi:10.1140/epje/i2010-10665-4

Foderà, V., van de Weert, M., & Vestergaard, B. (2010). Large-scale polymorphism and auto-catalytic effect in insulin fibrillogenesis. Soft Matter, 6(18), 4413–4419. doi:10.1039/C0SM00169D

Foderà, V., Vetri, V., Wind, T. S., Noppe, W., Cornett, C., Donald, A. M., … Vestergaard, B. (2014). Observation of the Early Structural Changes Leading to the Formation of Protein Superstructures. The Journal of Physical Chemistry Letters, 5(18), 3254–3258. doi:10.1021/jz501614e

Foderà, V., Zaccone, A., Lattuada, M., & Donald, A. M. (2013). Electrostatics Controls the Formation of Amyloid Superstructures in Protein Aggregation. Physical Review Letters, 111(10), 108105. doi:10.1103/PhysRevLett.111.108105

Galkin, O., & Vekilov, P. G. (2004). Mechanisms of Homogeneous Nucleation of Polymers of Sickle Cell Anemia Hemoglobin in Deoxy State. Journal of Molecular Biology, 336(1), 43–59. doi:https://doi.org/10.1016/j.jmb.2003.12.019

Garcia, G. A., Cohen, S. I. A., Dobson, C. M., & Knowles, T. P. J. (2014). Nucleation-conversion-polymerization reactions of biological macromolecules with prenucleation clusters. Physical Review E, 89(3), 032712. doi:10.1103/PhysRevE.89.032712

Goldenfeld, N. (1987). Theory of spherulitic crystallization. Journal of Crystal Growth, 84(4), 601–608. doi:https://doi.org/10.1016/0022-0248(87)90051-0

Gordon, M. P., Ha, T., & Selvin, P. R. (2004). Single-molecule high-resolution imaging with photobleaching. Proceedings of the National Academy of Sciences of the United States of America, 101(17), 6462–6465. doi:10.1073/pnas.0401638101

Gránásy, L., Pusztai, T., Tegze, G., Warren, J. A., & Douglas, J. F. (2005). Growth and form of spherulites. Physical Review E, 72(1), 011605. doi:10.1103/PhysRevE.72.011605

Hayashi, S., & Okada, Y. (2015). Ultrafast superresolution fluorescence imaging with spinning disk confocal microscope optics. Molecular Biology of the Cell, 26(9), 1743–1751. doi:10.1091/mbc.E14-08-1287

Heaney, P. J., & Davis, A. M. (1995). Observation and Origin of Self-Organized Textures in Agates. Science, 269(5230), 1562–1565. doi:10.1126/science.269.5230.1562

Hosier, I. L., Bassett, D. C., & Vaughan, A. S. (2000). Spherulitic Growth and Cellulation in Dilute Blends of Monodisperse Long n-Alkanes. Macromolecules, 33(23), 8781–8790. doi:10.1021/ma000946t

House, E., Jones, K., & Exley, C. (2011). Spherulites in Human Brain Tissue are Composed of Beta Sheets of Amyloid and Resemble Senile Plaques. Journal of Alzheimers Disease, 25(1), 43–46. doi:10.3233/jad-2011-110071

Huang, B., Jones, S. A., Brandenburg, B., & Zhuang, X. (2008). Whole-cell 3D STORM reveals interactions between cellular structures with nanometer-scale resolution. Nature Methods, 5, 1047. doi:10.1038/nmeth.1274

Huang, B., Wang, W., Bates, M., & Zhuang, X. (2008). Three-Dimensional Super-Resolution Imaging by Stochastic Optical Reconstruction Microscopy. Science, 319(5864), 810–813. doi:10.1126/science.1153529

Jensen, S. B., Thodberg, S., Parween, S., Moses, M. E., Hansen, C. C., Thomsen, J., … Hatzakis, N. S. (2021). Biased cytochrome P450-mediated metabolism via small-molecule ligands binding P450 oxidoreductase. Nature Communications, 12(1), 2260. doi:10.1038/s41467-021-22562-w

Jiang, Y., Shi, K., Xia, D., Wang, S., Song, T., & Cui, F. Protein Spherulites for Sustained Release of Interferon: Preparation, Characterization and <em>in vivo</em> Evaluation. Journal of Pharmaceutical Sciences, 100(5), 1913–1922. doi:10.1002/jps.22403

Kajioka, H., Hikosaka, M., Taguchi, K., & Toda, A. (2008). Branching and re-orientation of lamellar crystals in non-banded poly(butene-1) spherulites. Polymer, 49(6), 1685–1692. doi:https://doi.org/10.1016/j.polymer.2008.01.066

Krebs, M. R. H., Bromley, E. H. C., Rogers, S. S., & Donald, A. M. (2005). The mechanism of amyloid spherulite formation by bovine insulin. Biophysical Journal, 88(3), 2013–2021. doi:10.1529/biophysj.104.051896

Krebs, M. R. H., MacPhee, C. E., Miller, A. F., Dunlop, I. E., Dobson, C. M., & Donald, A. M. (2004). The formation of spherulites by amyloid fibrils of bovine insulin. Proceedings of the National Academy of Sciences of the United States of America, 101(40), 14420–14424. doi:10.1073/pnas.0405933101

Lu, Z. P., Goh, T. T., Li, Y., & Ng, S. C. (1999). Glass formation in La-based La–Al–Ni–Cu– (Co) alloys by Bridgman solidification and their glass forming ability. Acta Materialia, 47(7), 2215–2224. doi:https://doi.org/10.1016/S1359-6454(99)00058-0

Majumder, S., Busch, H., Poudel, P., Mecking, S., & Reiter, G. (2018). Growth Kinetics of Stacks of Lamellar Polymer Crystals. Macromolecules, 51(21), 8738–8745. doi:10.1021/acs.macromol.8b01765

Malle, M. G., Löffler, P. M. G., Bohr, S. S.-R., Sletfjerding, M. B., Risgaard, N. A., Jensen, S. B., … Hatzakis, N. S. (2021). Single particle combinatorial multiplexed liposome fusion mediated by DNA. bioRxiv, 2021.2001.2019.427313. doi:10.1101/2021.01.19.427313

Moses, M. E., Lund, P. M., Bohr, S. S. R., Iversen, J. F., Kæstel-Hansen, J., Kallenbach, A. S., … Hatzakis, N. S. (2021). Single-Molecule Study of Thermomyces lanuginosus Lipase in a Detergency Application System Reveals Diffusion Pattern Remodeling by Surfactants and Calcium. ACS Applied Materials and Interfaces, 13(28), 33704–33712. doi:10.1021/acsami.1c08809

Nielsen, L., Khurana, R., Coats, A., Frokjaer, S., Brange, J., Vyas, S., … Fink, A. L. (2001). Effect of Environmental Factors on the Kinetics of Insulin Fibril Formation: Elucidation of the Molecular Mechanism. Biochemistry, 40(20), 6036–6046. doi:10.1021/bi002555c

Ogi, H., Fukukshima, M., Hamada, H., Noi, K., Hirao, M., Yagi, H., & Goto, Y. (2014). Ultrafast propagation of β-amyloid fibrils in oligomeric cloud. Scientific Reports, 4(1), 6960. doi:10.1038/srep06960

Ovesný, M., Křížek, P., Borkovec, J., Švindrych, Z., & Hagen, G. M. (2014). ThunderSTORM: a comprehensive ImageJ plug-in for PALM and STORM data analysis and super-resolution imaging. Bioinformatics, 30(16), 2389–2390. doi:10.1093/bioinformatics/btu202

Pinholt, H. D., Bohr, S. S.-R., Iversen, J. F., Boomsma, W., & Hatzakis, N. S. (2021). Single-particle diffusional fingerprinting: A machine-learning framework for quantitative analysis of heterogeneous diffusion. Proceedings of the National Academy of Sciences, 118(31), e2104624118. doi:10.1073/pnas.2104624118

Pinotsi, D., Buell, A. K., Dobson, C. M., Schierle, G. S. K., & Kaminski, C. F. (2013). A Label-Free, Quantitative Assay of Amyloid Fibril Growth Based on Intrinsic Fluorescence. ChemBioChem, 14(7), 846–850. doi:doi:10.1002/cbic.201300103

Pinotsi, D., Buell, A. K., Galvagnion, C., Dobson, C. M., Kaminski Schierle, G. S., & Kaminski, C. F. (2014). Direct Observation of Heterogeneous Amyloid Fibril Growth Kinetics via Two-Color Super-Resolution Microscopy. Nano Letters, 14(1), 339–345. doi:10.1021/nl4041093

Qu, X., Wu, D., Mets, L., & Scherer, N. F. (2004). Nanometer-localized multiple single-molecule fluorescence microscopy. Proceedings of the National Academy of Sciences of the United States of America, 101(31), 11298–11303. doi:10.1073/pnas.0402155101

Ries, J., Udayar, V., Soragni, A., Hornemann, S., Nilsson, K. P. R., Riek, R., … Rajendran, L. (2013). Superresolution Imaging of Amyloid Fibrils with Binding-Activated Probes. ACS Chemical Neuroscience, 4(7), 1057–1061. doi:10.1021/cn400091m

Rogers, S. S., Krebs, M. R. H., Bromley, E. H. C., van der Linden, E., & Donald, A. M. (2006). Optical Microscopy of Growing Insulin Amyloid Spherulites on Surfaces In Vitro. Biophysical Journal, 90(3), 1043–1054. doi:10.1529/biophysj.105.072660

Sang, J. C., Hidari, E., Meisl, G., Ranasinghe, R. T., Spillantini, M. G., & Klenerman, D. (2021). Super-resolution imaging reveals α-synuclein seeded aggregation in SH-SY5Y cells. Communications Biology, 4(1), 613. doi:10.1038/s42003-021-02126-w

Shen, Y., Ruggeri, F. S., Vigolo, D., Kamada, A., Qamar, S., Levin, A., … Knowles, T. P. J. (2020). Biomolecular condensates undergo a generic shear-mediated liquid-to-solid transition. Nature Nanotechnology, 15(10), 841–847. doi:10.1038/s41565-020-0731-4

Shtukenberg, A. G., Punin, Y. O., Gunn, E., & Kahr, B. (2012). Spherulites. Chemical Reviews, 112(3), 1805–1838. doi:10.1021/cr200297f

Song, S., Zhou, H., Ye, S., Tam, J., Howe, J. Y., Manners, I., & Winnik, M. A. Spherulite-like Micelles. Angewandte Chemie International Edition, n/a(n/a). doi:https://doi.org/10.1002/anie.202101177

Stella, S., Mesa, P., Thomsen, J., Paul, B., Alcón, P., Jensen, S. B., … Montoya, G. (2018). Conformational Activation Promotes CRISPR-Cas12a Catalysis and Resetting of the Endonuclease Activity. Cell, 175(7), 1856-1871.e1821. doi:https://doi.org/10.1016/j.cell.2018.10.045

Tanaka, H., & Nishi, T. (1985). New Types of Phase Separation Behavior during the Crystallization Process in Polymer Blends with Phase Diagram. Physical Review Letters, 55(10), 1102–1105. doi:10.1103/PhysRevLett.55.1102

Thomsen, J., Sletfjerding, M., Jensen, S., Stella, S., Paul, B., Malle, M., … Hatzakis, N. (2020). DeepFRET, a software for rapid and automated single-molecule FRET data classification using deep learning. eLife, 9. doi:10.7554/eLife.60404

Thomsen, R. P., Malle, M. G., Okholm, A. H., Krishnan, S., Bohr, S. S. R., Sørensen, R. S., … Kjems, J. (2019). A large size-selective DNA nanopore with sensing applications. Nature Communications, 10(1), 5655. doi:10.1038/s41467-019-13284-1

Toprakcioglu, Z., Challa, P., Xu, C., & Knowles, T. P. J. (2019). Label-Free Analysis of Protein Aggregation and Phase Behavior. ACS Nano, 13(12), 13940–13948. doi:10.1021/acsnano.9b05552

Vetri, V., & Foderà, V. (2015). The route to protein aggregate superstructures: Particulates and amyloid-like spherulites. FEBS Letters, 589(19, Part A), 2448–2463. doi:http://doi.org/10.1016/j.febslet.2015.07.006

Yagi, H., Ban, T., Morigaki, K., Naiki, H., & Goto, Y. (2007). Visualization and Classification of Amyloid β Supramolecular Assemblies. Biochemistry, 46(51), 15009–15017. doi:10.1021/bi701842n

Zimmermann, M. R., Bera, S. C., Meisl, G., Dasadhikari, S., Ghosh, S., Linse, S., … Knowles, T. P. J. (2021). Mechanism of Secondary Nucleation at the Single Fibril Level from Direct Observations of Aβ42 Aggregation. Journal of the American Chemical Society, 143(40), 16621–16629. doi:10.1021/jacs.1c07228

