## Supplemental figures and table for "Direct Observation of Heterogeneous Formation of Amyloid Spherulites in Real-time by Super-resolution Microscopy"

**This PDF file includes:**

Figures S1 to S11
Tables S1
Legends for Movies S1 to S8

**Other supplementary materials for this manuscript include the following:**

Movies S1 to S8

### Supplementary Figures

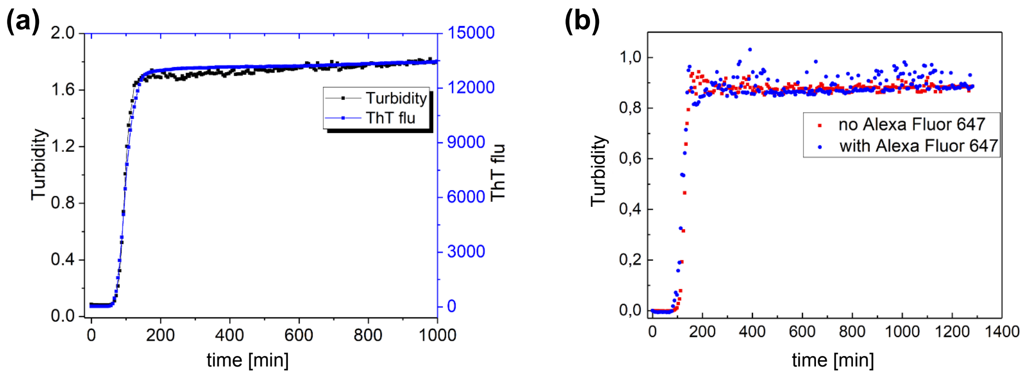


**Figure S1.** a) ThT fluorescence and turbidity kinetics showing that the aggregation process was complete in ~ 3 hours at 60 °C and that turbidity and fluorescence provided identical results, supporting amyloid origin. b) Turbidity kinetics of human insulin at 60 °C with and without Alexa Fluor 647. Each curve is the average on four replicates. The consistent of turbidities confirms that labeling doesn't interfere the aggregation process.

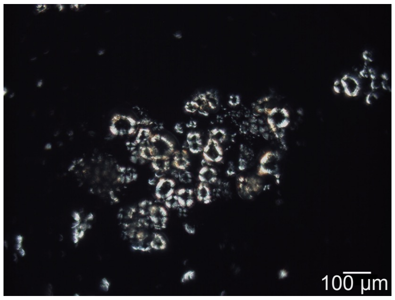


**Figure S2.** Spherulites images obtained by cross polarised microscopy Zeiss Axioplan Optical Microscope, Carl Zeiss. The characteristic Maltese cross shows spherulite formation. Scale bar is 100 μm.

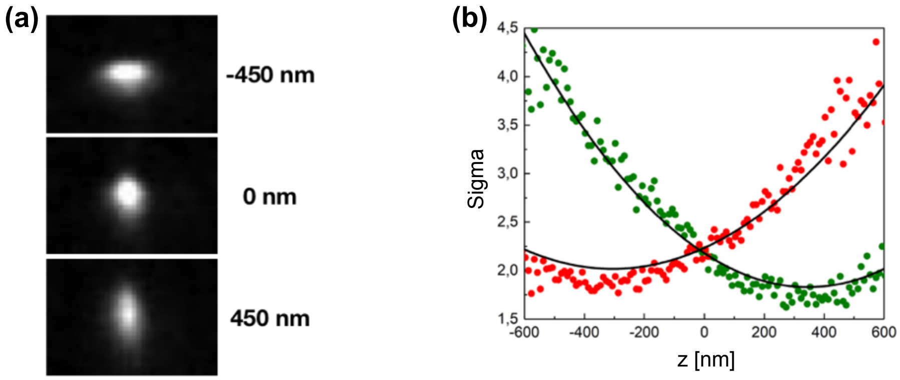


**Figure S3.** (a) Single localization imaging of Atto-655 labeled liposomes extruded at 50nm and tethered on poly-L-Lysine passivated surfaces. (b) Calibration curve of PSF widths as a function of z. Z step length was 10 nm. The calibration curve was calculated by ThunderSTORM.

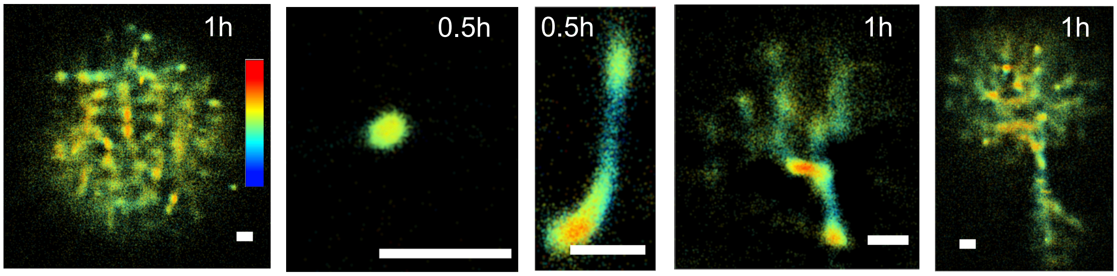


**Figure S4.** 3D dSTORM images at different steps of the aggregate growth at incubation time from 0.5 hour to 1 hours and incubation temperature at 60 °C. Pseudocolor scale is from −500 nm to 500 nm; Scale bars are 1 µm.

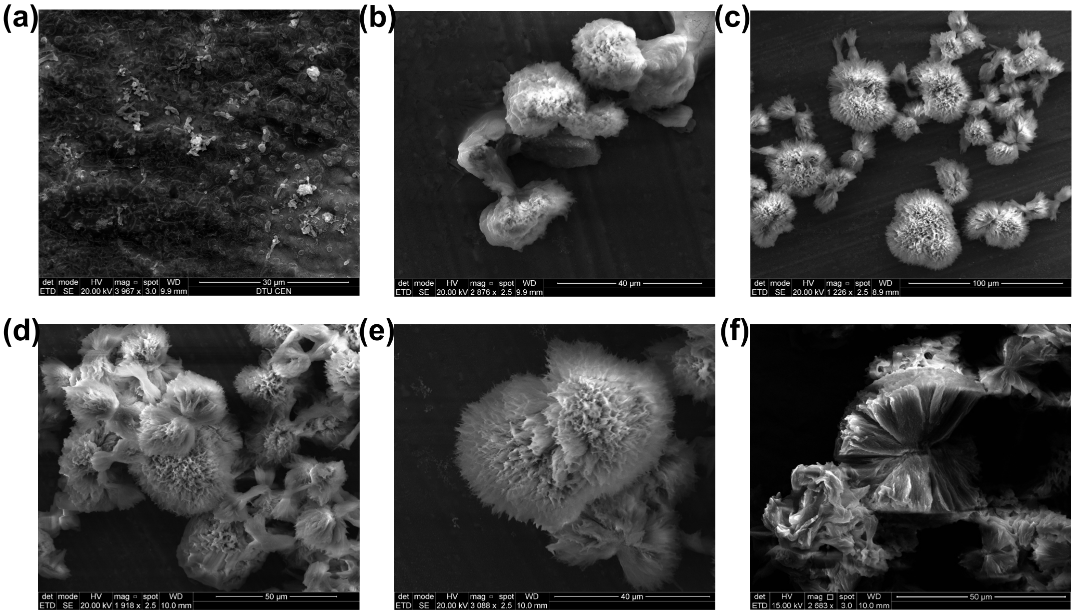


**Figure S5**. SEM image of spherulites grew at 60 ℃. (a) with incubation time of 1 hour in tube, (b) with incubation time of 2 hours in tube, (c) with incubation time of 4 hours and (d-e) with incubation time of 24 hours in tube. (e) a fully-grown insulin spherulite in tube, (f) with incubation time of 24 hours in microplate. Scale bars are (a) 30 µm, (b) 40 µm, (c) 100 µm, (d) 50 µm, (e) 40 µm and (f) 50 µm respectively.

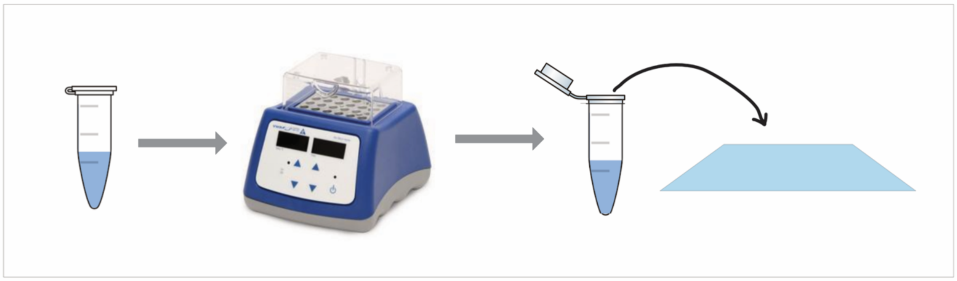

**Figure S6.** All the samples were pre-incubated in a block heater in the designated temperature to skip the lag-phase. Preincubated samples were then put on a microscope slide to monitor spherulite growth on the microscope.

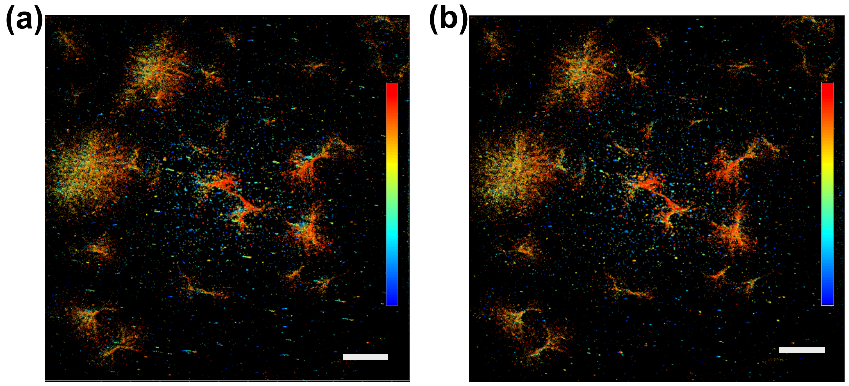

**Figure S7.** REPLOM image of HI spherulites grown at 45 °C (a) prior to and (b) after drift correction. Scale bars: 10 μm. The pseudocolor represents time and ranges from 0 s (blue) to 9500 s (red).

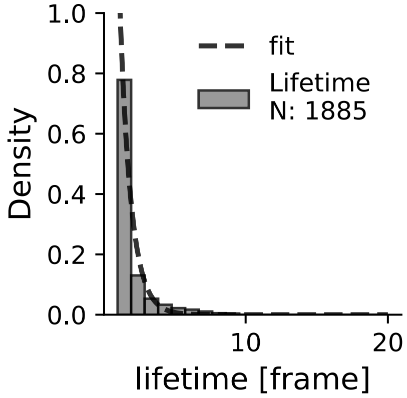


**Figure S8.** Quantification of bleaching time of Alexa 647 in REPLOM conditions. In the experimental condition used (absence of imaging buffer and high laser power) Alexa 647 chromophores are rapidly photobleached. Lifetime is 0.7845 ± 0.0017 frames.

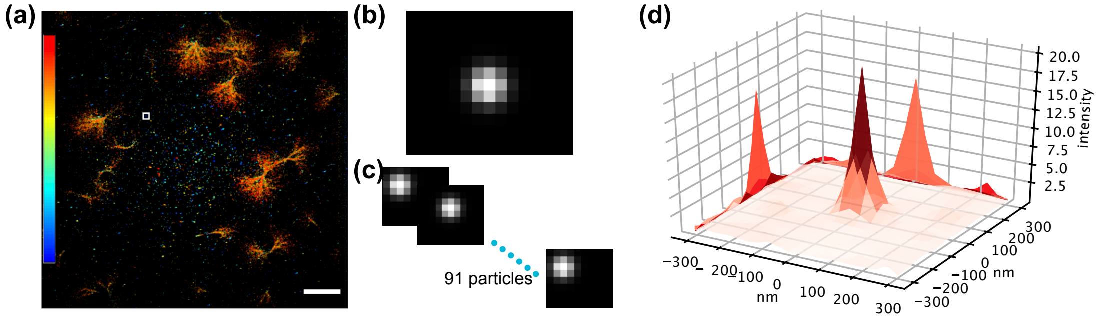


**Figure S9.** Quantification of REPLOM resolution. (a) REPLOM image of HI spherulites grew at 45 °C. Scale bar: 10 μm. (b) Higher-magnification view of the boxed region in (a), which contains a single spot. (c) 91 particles were selected for resolution calculation. (d) Resulting 2D histograms were generated by aligning the 91 particles to the same center. Two-dimensional gaussian fitting to the presented histogram allowed extraction of FWHM for resolution determination. FWHM x: 68.1 nm, FWHM y: 66.2 nm.

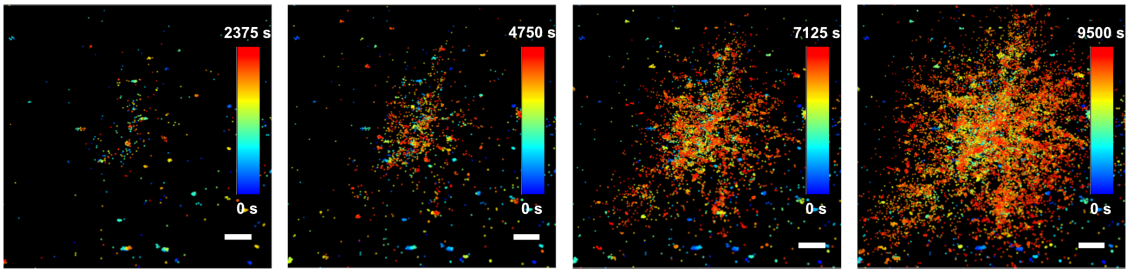


**Figure S10.** Direct real-time observation of temporal development of isotropic growth at t = 2375 time intervals. Scale bars: 2 μm.

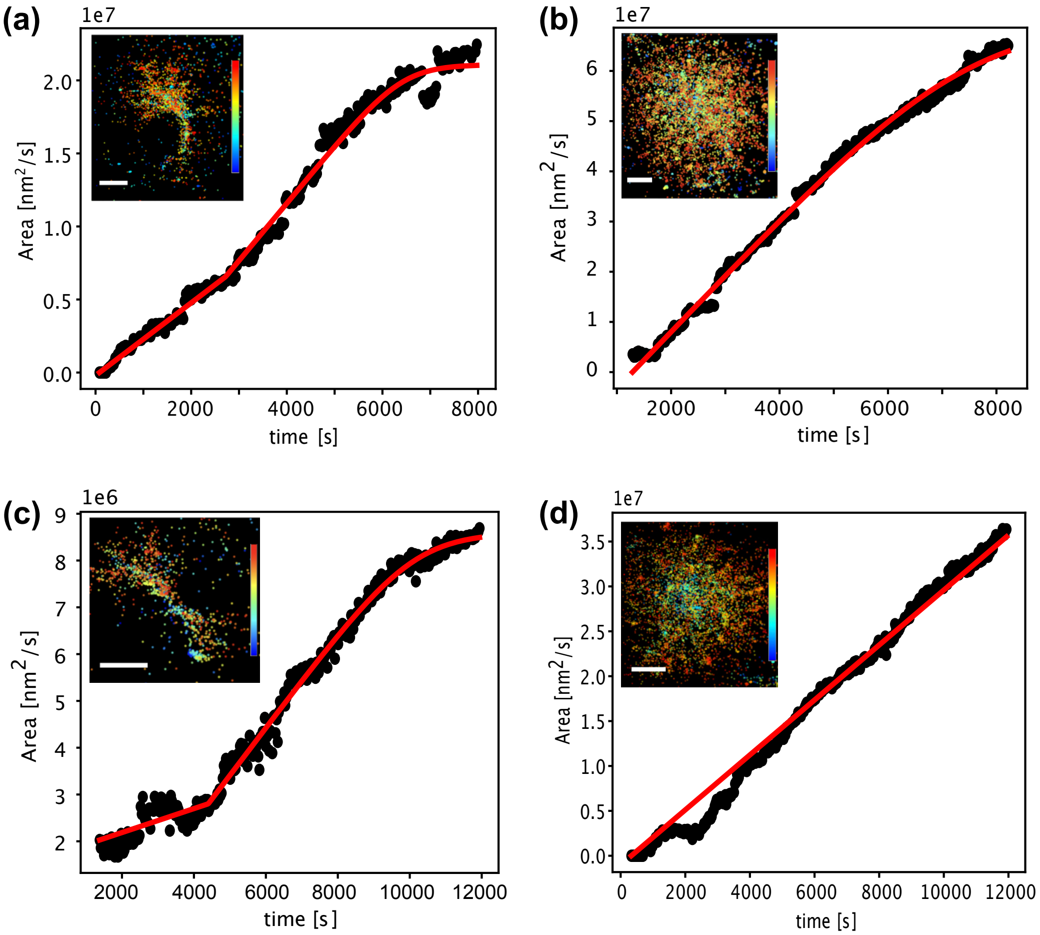


**Figure S11.** Representative anisotropic (a, c) and isotropic (b, d) HI spherulites obtained by REPLOM and their corresponding growth curves. The incubation temperatures were: (a&b) 37 °C, and (c&d) 32 °C. Scale bars: 2 um. In The pseudocolor bar represents time and spans from 0s to 8000s in (a and b) and from 0 s to 12000 s in (c and d).

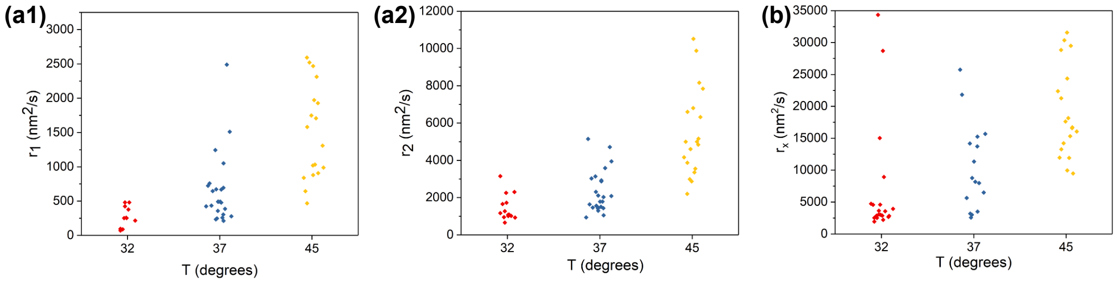


**Figure S12.** Growth rate distribution for all tested temperatures (32, 37 and 45 °C) for all spherulite growth morphologies. a1) for $r_1$ of anisotropic growth a2) for $r_2$ of anisotropic growth b) $r_x$ for isotropic growth. Data extracted from Arrhenius plots in Figure 4.

**Table S1.** Quantification of the diameters of the central fibrils of the intermediates in Fig 1.

| **Intermediates in Figure 1** | 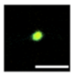 | 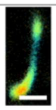 | 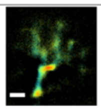 | 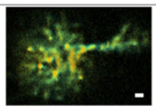 |
|---|---|---|---|---|
| **Diameter of the central fibrils (nm)** | ~300 | ~400 | ~450 | ~700 |

**Movie S1 (separate file).** 3D rotation video of anisotropic spherulite with asymmetric lobes.

**Movie S2 (separate file).** 3D rotation video of anisotropic spherulite with symmetric lobes.

**Movie S3 (separate file).** Real-time growth and the corresponding area of the anisotropic spherulite in Figure 3c (incubation temperature 45°C, exposure time of 30 ms followed by a waiting time for each frame of 25 s).

**Movie S4 (separate file).** Real-time growth and the corresponding area of the isotropic spherulite in Figure 3d (incubation temperature 45°C, exposure time of 30 ms followed by a waiting time for each frame of 25 s).

**Movie S5 (separate file).** Real-time growth and the corresponding area of the anisotropic spherulite in Figure S11a (incubation temperature 37°C, exposure time of 30 ms followed by a waiting time for each frame of 27.4 s).

**Movie S6 (separate file).** Real-time growth and the corresponding area of the isotropic spherulite in Figure S11b (incubation temperature 37°C, exposure time of 30 ms followed by a waiting time for each frame of 27.4 s).

**Movie S7 (separate file).** Real-time growth and the corresponding area of the anisotropic spherulite in Figure S11c (incubation temperature 32°C, exposure time of 30 ms followed by a waiting time for each frame of 30 s).

**Movie S8 (separate file).** Real-time growth and the corresponding area of the isotropic spherulite in Figure S11d (incubation temperature 37°C, exposure time of 30 ms followed by a waiting time for each frame of 30 s).
